# Differential Effects of Zooplankton on Sunlight Inactivation of Viruses

**DOI:** 10.64898/2026.03.05.709857

**Authors:** Martha I. Verbel-Olarte, Tamar Kohn, Niveen S. Ismail

**Affiliations:** Laboratory of Environmental Virology, School of Architecture, Civil and Environmental Engineering (ENAC), École Polytechnique Fédérale de Lausanne (EPFL), 1015 Lausanne, Switzerland; Picker Engineering Program, Smith College, Northampton, Massachusetts 01063, United States

**Keywords:** Viral inactivation, zooplankton grazing, solar disinfection, water treatment, microbial water quality

## Abstract

Interactions between viruses and filter-feeding zooplankton can alter viral persistence in surface waters, with direct implications for water quality and public health risk. However, data on virus-zooplankton interactions and the environmental factors that influence them are still limited. This study evaluated the impact of filter feeding, in the dark and under simulated sunlight, on a bacteriophage (MS2) and a human virus (echovirus11; E11) in the presence of a ciliate (*Tetrahymena pyriformis*) and rotifer (*Brachionus calyciflorus*). Dark experiments established organism-dependent baseline removal for each virus, and rotifers showed greater removal of both viruses in comparison to ciliates. Under simulated sunlight, in contrast, experiments with ciliates resulted in greater virus removal compared to experiments with rotifers over a similar timespan (4.2 vs. 2.7 log MS2 in 53-58 h; 3.5 vs. 3.0 log E11 in 24-25 h). Analysis of decay rate constants reveals species-specific shifts in virus removal between dark and light that, depending on viral type and zooplankton species, either accelerate viral attenuation or protect viruses and prolong infectivity. *T. pyriformis* increases removal under sunlight relative to dark conditions and acts synergistically with sunlight inactivation, whereas rotifers impede sunlight inactivation.

## 1. Introduction

Filter-feeding zooplankton regulate water quality by removing and ingesting suspended organic particles across a broad size range, from bacteria to algae. While their contribution to water quality improvement is well documented^1^, the interactions between zooplankton and microbial pollutants, particularly viruses, are complex and variable^2–7^. Zooplankton can reduce viral loads through predation and inactivate viruses during digestion, yet they can also harbor viruses^4,5,8–13^ and excrete infectious viruses in feces^14,15^, potentially serving as vectors. Viral inactivation is strongly dependent on environmental conditions^16^, and zooplankton-virus interactions vary with both zooplankton species and virus type^7,9,17–19^, but available data are insufficient to predict when grazing will contribute to viral decay or prolong infectivity. Understanding zooplankton-virus interactions and their influence on other environmental processes that modulate viral fate is essential for effective water quality management.

Interactions between zooplankton and viruses have direct consequences for human health and water safety. Enteric viruses in water bodies can cause illnesses such as gastroenteritis and respiratory infections^20,21^, and elevated microbial pollutant concentrations can prompt recreational water closures with substantial economic consequences^22^. To inform water quality management, research has characterized and modeled viral inactivation driven by abiotic factors, such as temperature and pH, with sunlight recognized as a major driver of viral inactivation^16,23– 28^. Environmental conditions and water chemistry, which modulate virus aggregation and surface adhesion, modify the efficacy of sunlight inactivation of viruses^23,29,30^. The impact of filter-feeding zooplankton on sunlight inactivation, however, remains inadequately characterized. Most prior studies addressed zooplankton effects on chemical and UV254 disinfection of bacteria^31–33^, or examined amoebae that use a predatory engulfing strategy rather than filter-feeding^4,9^. Numerous studies have demonstrated protective effects for bacteria within protists. For example, *Tetrahymena sp*. can protect coliform bacteria from chlorination^34^, and *Campylobacter* from virudine disinfectant^35^. One study observed protection of Φ174 and MS2 from UV by several *Tetrahymena* species^10^. For rotifers, studies have shown that they can harbor *Vibrio*, but protection effects specific to rotifers and bacteria or viruses are not well characterized^36^. To date, only one study has shown protection of MS2 from sunlight by the brackish-water rotifer, *Brachionus plicatilis*^37^.

In this study, we evaluate the effect of two filter-feeding zooplankton species, the ciliated protist *Tetrahymena pyriformis* and the rotifer *Brachionus calyciflorus*, on the decay of echovirus 11 (E11) and bacteriophage MS2 in the presence and absence of sunlight. E11, an enterovirus associated with a broad spectrum of human illnesses^38^, is commonly detected in environmental waters and wastewater ^39^, and has been shown to be removed by filter-feeding zooplankton ^7,12^. MS2 is widely used as a surrogate for human enteric viruses^18^ and is considered a conservative indicator of sunlight inactivation of these viruses. Its removal has also been investigated in co-incubation studies with filter-feeding zooplankton^7,18^. Studying these two viruses enables comparison with prior work while providing new insights into zooplankton-mediated effects on their environmental persistence.

The two zooplankton species were chosen for their relevance to both natural freshwater environments and engineered treatment systems, as well as their extensive use as model organisms^40,41^. *T. pyriformis* has been widely adopted as a model species, yet prior research has reported substantial variability in viral uptake, even when examining the same virus across different studies ^7,12,13,42^. In contrast, viral interactions with *B. calyciflorus* remain relatively understudied, providing an opportunity to extend understanding beyond the limited data available for the brackish-water rotifer *B. plicatilis*. Given the variability in findings on viral-zooplankton dynamics and the scarcity of research on their role in sunlight inactivation of viruses, this study aims to quantify how dark and sunlight inactivation rates vary among viruses and zooplankton species within a single system.

## 2. Materials and Methods

### 2.1 *Tetrahymena pyriformis* Culture and Preparation

Cultures of *T. pyriformis* ciliates (Culture Collection of Algae and Protozoa 1630/1W) were maintained axenically in proteose peptone yeast extract (PPYE) medium at 24°C in 150 cm^2^ cell culture flasks (TTP, Milian). PPYE was prepared by dissolving 20.0 g of proteose peptone (VWR) and 2.5 g of yeast extract (Thermo Scientific) in 1 L of MilliQ water. The solution was autoclaved and stored at 4 °C. Subcultures were axenically prepared weekly in a laminar flow hood by spiking 1 mL of the previous culture into 39 mL of fresh PPYE (total volume of 40 mL). One day prior to the experiments, ciliates were washed according to previously published protocols ^3,7,12^. Briefly, 120 mL of *T. pyriformis* in PPYE were centrifuged at 400x*g* for 10 min to form a pellet. The supernatant was removed, and 10 mL of sterile phosphate-buffered saline (PBS) (Table S1) was added. The pellet was gently resuspended and centrifuged at 400x*g* for 6 min. The resuspension procedure was repeated twice with PBS, and the final pellet was suspended in 50 ml of filter-sterilized, moderately hard synthetic freshwater (MHSFW) (Table S2)^43^. The *T. pyriformis* suspension was left at room temperature for approximately 20 hours before use. *T. pyriformis* were enumerated using an automated cell counter in brightfield mode (DeNovix, CellDrop).

### 2.2 *Brachionus calyciflorus* Culture and Preparation

*B. calyciflorus* resting cysts (Florida Aquafarms) were hatched in 30 mL of sterile MHSFW and incubated at 25°C under low light intensity (17 μmol photons m^-2^ s^-1^) in an incubator (AlgaeTron AG 230) on a 16h:8h light: dark cycle. Upon hatching, rotifers (20-50 individuals) were transferred into a 1 L glass bottle containing 900 mL of MHSFW with high aeration and fed daily with 25 mL of the algae *Chlorella vulgaris* (SI text for culture details and Table S3). Cultures were refreshed weekly; one-half of the culture was discarded, and aggregates of algae and dead rotifers were removed by sieving through a 125 μm sieve. The collected filtrate from the 125 μm was subsequently sieved through a 50 μm sieve to collect the rotifers. The rotifers were removed from the sieve and resuspended in 900 mL of MHSFW. Two hours prior to each experiment, rotifers were concentrated using a 50 μm sieve and resuspended to the desired experimental density. Rotifers were enumerated using a gridded Sedgewick Rafter (Wildco) using an inverted light microscope (Olympus CK X41).

### 2.3 Virus Propagation and Preparation

Echovirus 11 (E11) (Gregory strain, ATCC^®^ VR37™) was propagated on Buffalo green monkey kidney (BGMK) cells (kindly provided by the Spiez Laboratory, Switzerland) using a previously published protocol^44^. E11 stock solutions were purified using 100 kDa centrifugal filters (Centricon), then stored at −20 °C and thawed immediately before use. Infectious E11 concentrations were determined by the most probable number (MPN) assay using serial 10-fold dilutions (20 μL inoculum) on 95% confluent BMGK cells incubated for 5 days following previously published methods^44^. E11 concentrations were reported as most probable number or cytopathic units per milliliter (MPNCU mL^-1^). RStudio software (Version 2023.06.1+524) was used to calculate the E11 MPNCU mL^-1^ from the infectivity assay. The limit of quantification (LOQ) of the MPN assay is 30 MPNCU mL^-1^, corresponding to the lowest E11 concentration with a 95% probability of detecting at least one well with a positive cytopathic effect under the experimental conditions.

MS2 (DSM 13767) bacteriophage was replicated in *Escherichia coli* (DSM 5695) host. Phage purification and propagation followed previously published protocols, with slight modifications ^45^. Briefly, bacterial cell debris was removed by centrifugation at 4000x*g* for 15 min, followed by vacuum filtration through a 0.45 μm PVDF membrane (Stericup) under sterile conditions. The MS2 was further purified using 100 kDa centrifugal filters by transferring 65 mL of phage solution to the filters and centrifuging at 3000x*g* for 30 min. An equal volume of PBS (65 mL) was added, followed by centrifugation. The purified viral stock solution was then eluted at 3000x*g* for 2 min and stored at −20 °C. Infectious MS2 (100 μL samples) was enumerated by the double-layer agar method, and concentrations are reported as plaque-forming units per mL (PFU mL^-1^)^37,46^. The LOQ for MS2 is 300 PFU mL^-1^.

### 2.4 Virus and Zooplankton Co-incubation Experiments

#### Sunlight Exposure Experiments

Simulated sunlight exposure experiments were conducted using a solar simulator (Sun 2000; Abet Technologies) equipped with a 1,000-W Xenon lamp, an Air Mass (AM) 1.5 filter, and a 2-mm thick atmospheric edge (AE) filter, and was operated at 30 A. The irradiance spectrum was measured at four locations below the simulator using a spectroradiometer (ILT 900-R; International Light Technologies, Peabody, MA) (Figure S1). The average UVB irradiance, calculated over the wavelength range from 280 to 315 nm, was 0.17 ± 0.005 W m^-2^. This irradiance was selected to preserve healthy zooplankton populations while still allowing measurable sunlight inactivation kinetics. Experimental vessels were 400 mL glass beakers painted black to prevent sunlight from entering through the sides. Beakers were placed in a recirculating water bath to maintain a temperature of 24°C.

Experimental treatments (n=6 per virus and zooplankton combination) contained zooplankton (*T. pyriformis* or *B. calyciflorus*) in 150 mL MHSFW (water depth, z=3.5 cm) co-incubated with virus to achieve initial concentrations of 10^6^ PFU mL^-1^ for MS2 and 10^5^ MPNCU mL^-1^ for E11. Constant aeration was provided to maintain mixing and sufficient dissolved oxygen levels. The average initial experimental *T. pyriformis* density was 10^5^ ciliates mL^-1^ and the average initial density for *B. calyciflorus*, was 130 rotifers mL^-1^. Zooplankton viability and densities were checked at the start, midpoint, and end of each experiment. Concentrations varied by less than 10% for rotifers (Figure S2) and less than 15% for ciliates (Figure S3) over each experiment. MS2 light exposures lasted 53-58 hours, and the E11 exposures lasted 24-25 hours. Samples were collected periodically, filtered through a 0.22 μm sterile syringe filter (FILTER-BIO) to remove zooplankton, and infectious virus concentrations were enumerated. Unfiltered aliquots were utilized for UV-vis absorbance measurements (see below), zooplankton enumeration, and assessment of zooplankton viability through visual inspection using an inverted light microscope. Virus-only controls (n=2) were run in sterile MHSFW in parallel to quantify viral sunlight inactivation in the absence of zooplankton.

#### Dark Experiments

Dark experiments with zooplankton (n=6) and controls without zooplankton (n=3) followed the same protocol as the sunlight-exposure experiments, except that the beaker tops were covered with aluminum foil to exclude light. Dark experiments quantified dark biotic removal of viruses and baseline dark decay due to abiotic losses. Incubation durations were chosen based on the duration of the sunlight inactivation experiments. For MS2, dark experiments ran for 55 hours, and for E11, exposure ran for 30 hours.

### 2.5 Data analysis

The irradiance values were corrected for light absorbance by solutions containing rotifers and ciliates using a correction (screening) factor (Equation 1)^23,47^. The absorbance spectrum (α_s_) of each solution was obtained using a UV-vis with an integrating sphere (Shimadzu, model UV 2450–2550) to account for light scattering by the solution (Figure S4).

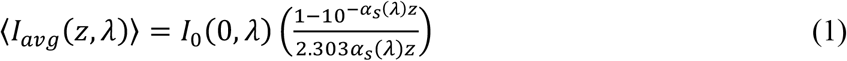

where ⟨*I*_*avg*_(*z,λ*)⟩ in W cm^-2^ nm^-1^ is the spectral irradiance averaged over the depth *z* of the experimental solution in each beaker in cm, *I*_*0*_(*0,λ*) is the incident spectral irradiance in W cm^-2^ nm^-1^, and α_s_ (*λ*) is the absorbance spectrum of the experimental solutions in cm^-1^.

Rate constants were calculated with respect to time (*k*, hr^-1^) and fluence (*κ*, m^2^ kJ^-1^) for pooled samples across all experimental replicates using a first-order decay model:

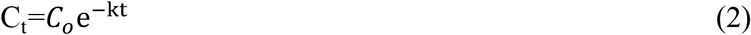

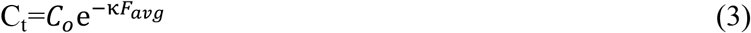

where *C*_*o*_ is the virus concentration at t = 0 hours in PFU mL^-1^ for MS2 or MPNCU mL^-1^for E11, *C*_*t*_ is the virus concentration at a given time point, *t* is time in hours, and *F*_*avg*_ is the UVB fluence averaged over the depth *z* of the experimental solution in each beaker in kJ m^-2^ calculated as:

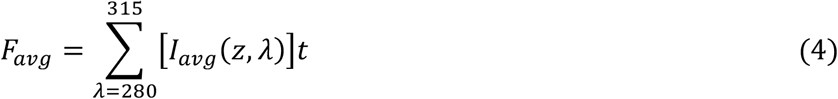

*F*_*avg*_ is referred to as fluence throughout the manuscript.

All statistical analyses were completed using SPSS (v28, IBM). All viral concentration data were log-transformed before statistical analysis. Results were considered significant for p<0.05. The mean ± standard deviation or propagated error values are presented for experimental data. Both standard deviation and propagated error are referred to as SD in the manuscript. The standard error (SE) is reported for calculated rate constants (*k*-value and *κ*-values). Repeated-measures ANOVA and linear mixed models (LMM) were used to test for differences in viral concentration across treatments in the experiments.

## 3. Results and discussion

### 3.1 Virus type and zooplankton species impact the effectiveness of dark inactivation

The removal rates of MS2 and E11 were first quantified in the dark, in the presence of *T. pyriformis* and *B. calyciflorus*, with appropriate controls (Figures 1A and 1B). Control beakers kept in the dark without zooplankton (virus-only controls) showed an average 0.5 ± 0.1 log decline over the experimental time period for both MS2 (55 h) and E11 (30 h), indicating that abiotic factors under dark conditions do not cause substantial inactivation. This observation is consistent with previous reports of enteric virus and phage stability for days, depending on the system conditions and the water matrix^2^.

**Figure 1.**
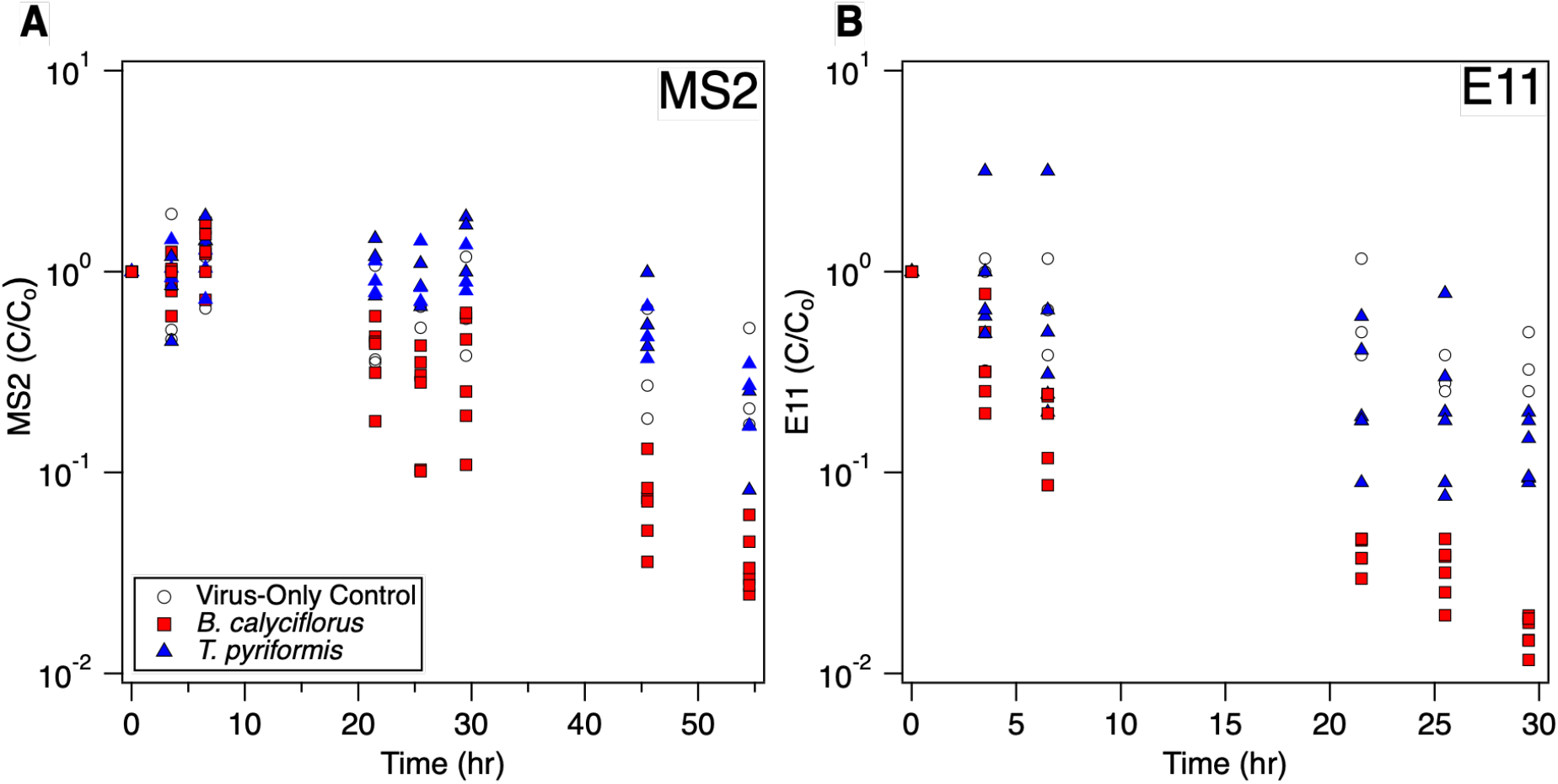
Viral decay in controls without zooplankton and removal kinetics during co-incubation with zooplankton in the dark. A) MS2 and B) E11. Six replicates were tested with zooplankton, and virus-only controls without zooplankton were tested in triplicate. Note the differing time scales on the x-axes for the two viruses.

When MS2 was incubated with *T. pyriformis*, decay was not significantly different from the virus-only control (ANOVA, p=0.180, Figure 1A). After 55 hours, treatments with ciliates showed a 0.7 ± 0.1 log removal, while virus-only controls showed a 0.6± 0.1 log decay. Previous studies reported higher MS2 removal by *T. pyriformis*: 1.3 log removal in ciliate treatments versus 0.6 log decay in virus-only controls over 48 hours in MHSFW^7^, and about 1.1 log removal with ciliates and 0.5 log decay in virus-only controls after 48 hours in a salt solution^18^. Although these values are numerically higher than ours, when considering net log removal (treatment minus control), all reported removals are less than 1 log and thus likely of limited biological relevance. In contrast, *B. calyciflorus* produced a 1.5 ± 0.01 log removal of MS2 in the dark compared with a 0.6± 0.1 log reduction in the virus-only controls. Across the full time series, the decline in virus concentration in the rotifer treatments was significantly greater than in the virus-only controls (ANOVA, p=0.032, Figure 1A). Comparative data for rotifers are limited, but *B. plicatilis*, a brackish-water rotifer of the same genus with similar feeding behavior, achieved a net 2 to 3 log removal of MS2 (after accounting for decay in the control) over 96 hours in brackish water^37^.

Similar findings were obtained for E11. Incubation with *T. pyriformis* did not significantly increase dark removal relative to the control with 0.9 ± 0.1 log removal over 30 hours with ciliates and 0.5 ± 0.1 log decay in virus-only controls (ANOVA, p=0.258, Figure 1B). Previous work has shown that E11 removal by *T. pyriformis* can require up to 96 hours to achieve 1.5 log removal, with only 0.3 log removal at 48 hours ^12^. In contrast, *B. calyciflorus* significantly enhanced E11 removal in the dark, achieving 1.8 ± 0.01 log removal versus a 0.5 ± 0.1 log decay in virus-only controls (ANOVA, p<0.001, Figure 1B). Although *B. calyciflorus* removed both MS2 and E11, the MS2-rotifer system required 55 hours to reach 1.5 log decline, whereas E11 reached 1.8 log decline within 30 hours under the same dark conditions.

The dark experiments established baseline information on virus stability in the water matrix used herein and showed differences in removal as a function of zooplankton species and virus. Previous research has shown variability in viral uptake by *T. pyriformis*^7,10,18^, but the underlying causes remain unclear. For example, Olive et al. found that uptake is virus-specific and may be linked to virus surface hydrophobicity, but not to virus size ^7^. In our study, E11 and MS2 are similarly sized, yet significant differences in removal were observed (LMM, p<0.001), consistent with the findings of Olive et al^7^.

Mechanistic data on rotifer uptake of viruses are limited beyond MS2 removal by *B. plicatilis*^37^. Studies of other particle types indicate that *B. calyciflorus* may alter feeding behavior according to particle characteristics, concentration, and size^48^. Rotifers can ingest particles in the bacterial size range, but their ciliary feeding structures are not optimized to collect particles in the sub-micron range^48,49^. In our experiments, rotifers removed viruses in the 27-30 nm range, likely at reduced clearance rates compared with bacteria.

Given the zooplankton densities used in this study and published clearance rate ranges, both *T. pyriformis* and *B. calyciflorus* would be expected to have similar theoretical capacities to process the beaker contents. Reported clearance rates for *T. pyriformis* on 1μm microspheres (surrogates for bacteria) are 5x10^-5^ to 1x10^-4^ mL ciliate^-1^ hr^-1 50^, which at our density (10^5^ ciliates mL^-1^) corresponds to roughly 130-250 passes of the beaker over 25 hours. Under other conditions, *T. pyriformis* clearance of E11 has been estimated at 1x10^-7^ mL ciliate^-1^ hr^-1^, corresponding to 0.25 passages over 25 hours^7^. For *B. calyciflorus*, reported bacterial clearance rates range from 1x10^-4^ to 5x10^-2^ mL rotifer^-1^ hr^-1, 48^ corresponding to 0.33 to 160 passes of the beaker over 25 hours using our experimental density of 130 rotifers mL^-1^. Although these published values are not specific to our exact conditions, they suggest comparable theoretical capacities for water processing by both zooplankton species at the densities used. However, we observed significantly different removal between *T. pyriformis* and *B. calyciflorus* for both MS2 and E11 (ANOVA, p<0.001), underscoring the biological complexity and the need to further elucidate the mechanisms driving these differences.

### 3.2 Sunlight inactivation of viruses is variably impacted by ciliates and rotifers

Figures 2A and 2B compare MS2 removal by simulated sunlight in the absence and presence of *T. pyriformis*. Time-based analysis (Figure 2A) suggests differences in decay rates of MS2 with and without *T. pyriformis*. However, after correcting for absorbance (Eq. 1, Figure S4), treatments with *T. pyriformis* exhibited a steeper decline than the controls. Sunlight alone achieved a total decay of approximately 2.5 log at a fluence of 32 kJ m^-2^, whereas the presence of *T. pyriformis* significantly enhanced MS2 decay to approximately 4.2 log at a fluence of only 17 kJ m^-2^ (LMM, p=0.029). Note that the strong absorbance of solutions containing *T. pyriformis* (Figure S4) caused the pronounced reduction in the fluence in *T. pyriformis* experiments compared to the virus-only controls, despite equal experimental times. For *B. calyciflorus* experiments (Figures 2C & 2D), sunlight alone resulted in approximately 2.0 log decay at a fluence of 29 kJ m^-2^ compared with 2.7 log decay at a fluence of 31 kJ m^-2^. Although there was a 0.7 log difference between the control and experimental treatments at the end point, this difference was not statistically significant when considering the entire time series (LMM, p=0.174) or fluence data (LMM, p=0.171).

**Figure 2.**
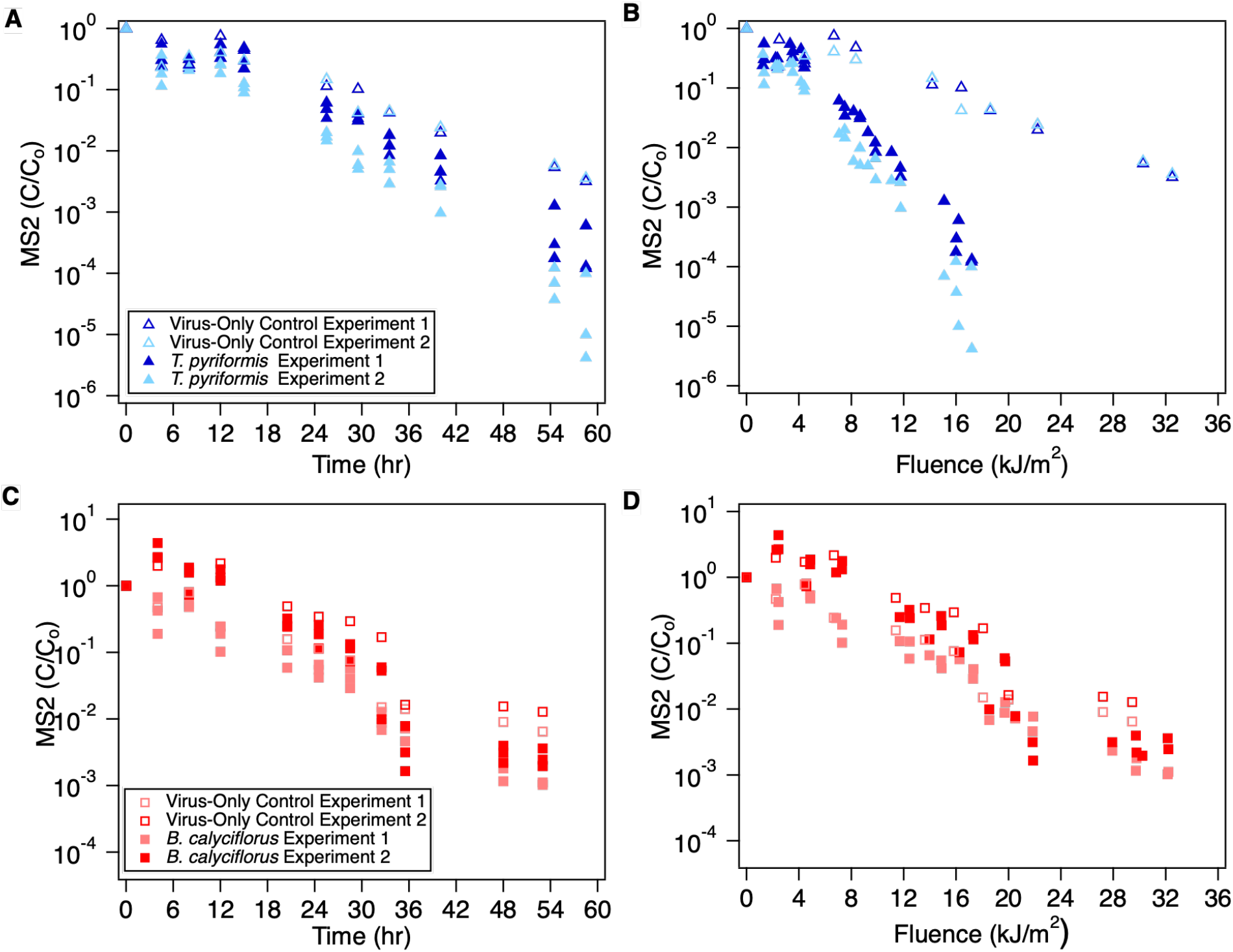
Sunlight inactivation of MS2 in the absence and presence of zooplankton. Panels A and B show the effect of *T. pyriformis* on MS2 concentration as a function of time and fluence, respectively. Panels C and D show the impact of *B. calyciflorus* as a function of time and fluence, respectively. Data from three replicates are presented for each of the zooplankton experiments (n=3). Note the difference in y-axis values for *T. pyriformis* (A and B) versus *B. calyciflorus* (C and D).

*T. pyriformis* also significantly enhanced E11 removal under sunlight. An enhancement was observed when E11 decay was analyzed as a function of time (Figure 3A) (LMM, p=0.044), and it was even more pronounced when analyzed as a function of fluence, with a 3.5 log decay at 7.3 kJ m^-2^ fluence in the presence of ciliates, compared with 2.2 log decay at 14.1 kJ m^-2^ fluence in the virus-only controls (LMM, p=0.002) (Figure 3B). In contrast, *B. calyciflorus* significantly protected E11 from sunlight inactivation. This effect, too, was evident both as a function of time (LMM, p <0.001) and fluence ( LMM, p<0.001) (Figures 3C & D), with 3.0 log decay at 14.5 kJ m^-2^ fluence with rotifers versus 4.0 log decay at 11.2 kJ m^-2^ fluence for the virus-only controls. Virus-only controls exhibited variable E11 behavior across experiments, which may be due to the sensitivity of E11 to aeration. Aeration sensitivity was not observed for MS2. Previous studies have reported virus-specific sensitivity to aeration. For example, Qβ was sensitive to aeration, while MS2 and ΦX174 were not^51^. In our experiments, vigorous aeration was particularly necessary for rotifer experiments to provide sufficient dissolved oxygen, whereas for ciliates, aeration was provided to ensure adequate mixing without causing cyst formation or death. Because a simple aquarium air pump was used, air flow rates were not measured.

**Figure 3.**
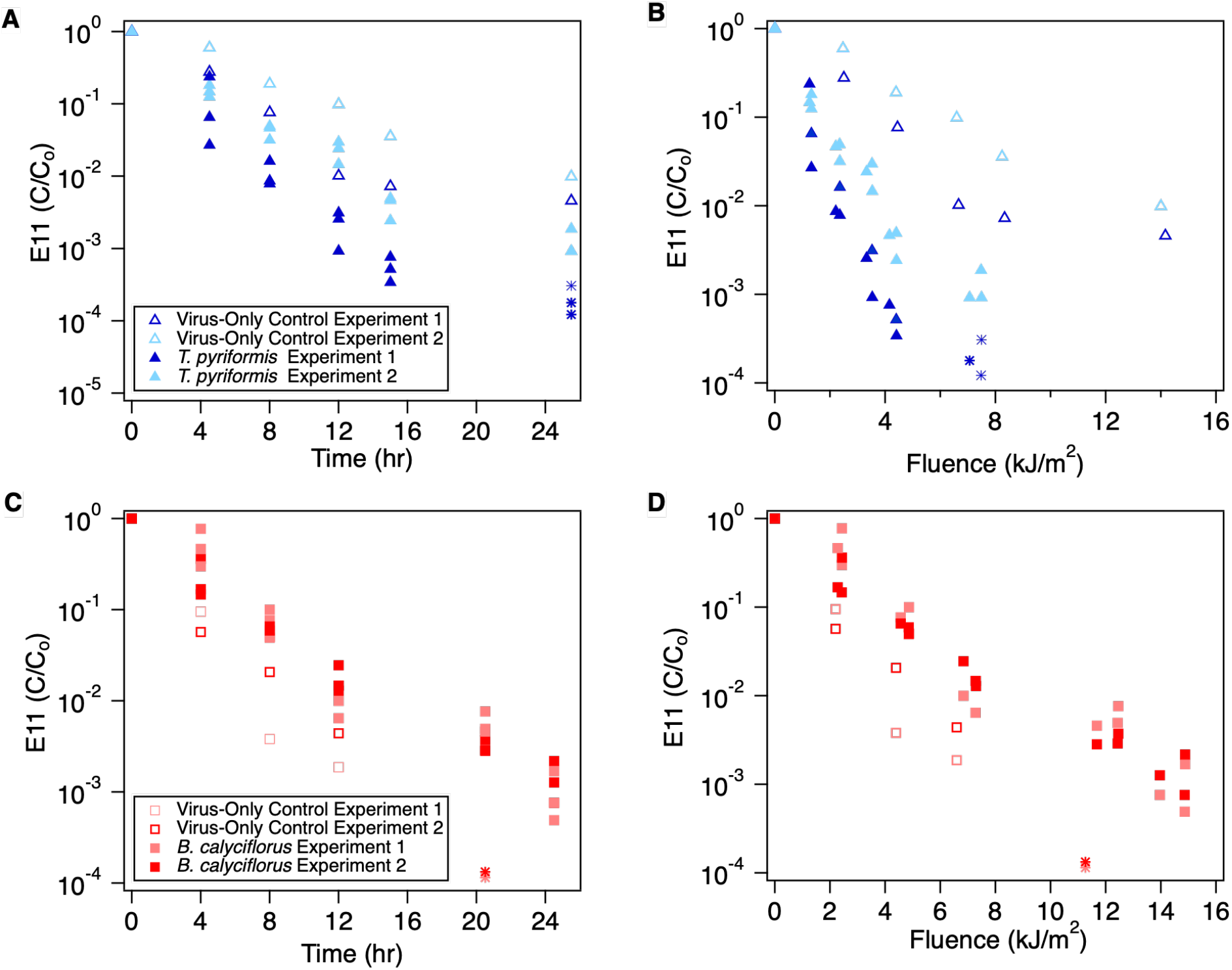
Sunlight inactivation of E11 in the absence and presence of zooplankton. Panels A and B show the effect of *T. pyriformis* on E11 concentration as a function of time and fluence, respectively. Panels C and D show the impact of *B. calyciflorus* as a function of time and fluence, respectively. Data from three replicates are presented for each of the zooplankton experiments (n=3). Star symbols indicate infectious virus concentrations below the limit of quantification.

Limited knowledge exists regarding how rotifers and ciliates affect the inactivation of viruses by sunlight. One study reported modest protection (3-8% reduction in inactivation ) of Φ174 and MS2 by several *Tetrahymena* species, including *T. pyriformis*, during a 3-minute exposure to UVB light^10^. Our results contrast with these previous findings, as decay of both E11 and MS2 under sunlight in the UVB range was enhanced in the presence of *T. pyriformis*. Differences in system conditions, including media type and exposure conditions, may explain the conflicting results. For rotifers, protection of MS2 from sunlight by *B. plicatilis* in brackish water has been reported^37^. In our system, *B. calyciflorus* strongly protected E11, but effects on MS2 were minor and not statistically significant. Differences in media, rotifer species, and experimental conditions limit direct comparison of MS2 data across studies.

### 3.3 Comparison of inactivation rate constants reveals synergistic effect of *T. pyriformis* and antagonistic effects of *B. calyciflorus* on sunlight inactivation

To further quantify zooplankton impacts on virus inactivation under dark and sunlight conditions, we calculated inactivation rate constants with respect to time (k, hr^-1^) and fluence (κ, m^2^ kJ^-1^) (Table 1). Because solution absorbance differed between treatments, fluence-based κ-values are the most appropriate metric for comparing sunlight experiments. Experimental to virus-only control ratios for time-based *k* (dark) and fluence-based *κ* (light) were calculated to facilitate comparisons of zooplankton effects in dark and light conditions (Table S4). For example, for E11 with *T. pyriformis*, the dark *k*-ratio (experimental/virus-only control) was 1.8 while the light *κ*-ratio was 3.1, suggesting an approximately 1.7 fold enhancement of removal by *T. pyriformis* in the light. A similar pattern was observed for MS2 with *T. pyriformis*: the dark *k*-ratio was 0.66, but as previously noted, there was no significant difference in dark experimental (with *T. pyriformis*) and dark control (virus-only) time series data. In the light for *T. pyriformis*, the *κ*-ratio was 3.2 for MS2. Combined, these analyses indicate altered feeding or processing behavior of *T. pyriformis* in sunlight in comparison to the dark. By contrast, *B. calyciflorus* showed greater removal than the virus-only control in the dark for E11 with a *k*-ratio of 3.8, but they produced a fractional *κ*-ratio of 0.57 in the light, indicating protection of E11 from the disinfection effects of sunlight. While a similar trend in the ratios is also observed for *B. calyciflorus* co-incubation with MS2, as previously discussed, differences between the control and experimental time-series and fluence-based data under sunlight were not statistically significant.

**Table 1.** Virus inactivation rate constants based on time (k, hr^-1^) and fluence ( κ, m^2^ kJ ^−1^) under light and dark conditions, with and without zooplankton (virus-only control). Standard errors are reported for each rate constant

| Condition | *T. pyriformis*<br>Experimental | Virus-Only<br>*T. pyriformis* Control | *B. calyciflorus*<br>Experimental | Virus-Only<br>*B. calyciflorus* Control |
| --- | --- | --- | --- | --- |
| Dark E11 | 0.068±0.008 $\text{hr}^{-1}$ | 0.038±0.005 $\text{hr}^{-1}$ | 0.14±0.006 $\text{hr}^{-1}$ | 0.038±0.005 $\text{hr}^{-1}$ |
| Light E11 | 0.37±0.02 $\text{hr}^{-1}$ | 0.23±0.02 $\text{hr}^{-1}$ | 0.29±0.007 $\text{hr}^{-1}$ | 0.47±0.02 $\text{hr}^{-1}$ |
| | 1.3±0.06 $\text{m}^2 \text{kJ}^{-1}$ | 0.42±0.03 $\text{m}^2 \text{kJ}^{-1}$ | 0.49±0.01 $\text{m}^2 \text{kJ}^{-1}$ | 0.86±0.04 $\text{m}^2 \text{kJ}^{-1}$ |
| Dark MS2 | 0.016±0.005 $\text{hr}^{-1}$ | 0.024±0.002 $\text{hr}^{-1}$ | 0.056±0.005 $\text{hr}^{-1}$ | 0.024±0.002 $\text{hr}^{-1}$ |
| Light MS2 | 0.15±0.004 $\text{hr}^{-1}$ | 0.094±0.003 $\text{hr}^{-1}$ | 0.12±0.005 $\text{hr}^{-1}$ | 0.088±0.006 $\text{hr}^{-1}$ |
| | 0.54±0.01 $\text{m}^2 \text{kJ}^{-1}$ | 0.17±0.006 $\text{m}^2 \text{kJ}^{-1}$ | 0.20±0.008 $\text{m}^2 \text{kJ}^{-1}$ | 0.16±0.01 $\text{m}^2 \text{kJ}^{-1}$ |

Additive effects for *T. pyriformis* were assessed by comparing the sum of dark inactivation by *T. pyriformis* (*k*_*Dark, T. pyriformis*_) and sunlight inactivation for the virus-only control (*k*_*Light, virus-only control*_) with the experimental sunlight inactivation with *T. pyriformis* (*k*_*Light, T. pyriformis*_)(Table S5). For both MS2 and E11, the summed time-based rate constants were lower than the observed *k*_*Light, T. pyriformis*_; for MS2, the values were 0.11 versus 0.15, and for E11, 0.30 versus 0.37, indicating synergistic enhancement of inactivation when grazing and sunlight act together. Because time-based inactivation rates do not account for differences in solution absorbance and delivered fluence, these comparisons likely underestimate the synergistic effect.

Differences in decay rate constant ratios between dark and light conditions may reflect behavioral and physiological changes in zooplankton. Phototactic responses differ among zooplankton. *Brachionus spp*. possess an eye spot that senses light, and exhibit phototaxis that varies with light intensity and wavelength^52^. Rotifers show positive phototaxis at different wavelengths in the visible light spectrum and negative phototaxis in UVA light^53^, but UVB light has not examined. Although *T. pyriformis* does not have known photoreceptors, it has exhibited negative phototaxis under a high intensity mercury vapor light at certain tested wavelengths from 250-700 nm^54^. This response may be driven by heat, or *Tetrahymena spp*. may contain rhodopsin-like proteins involved in a light response^55^.

Beyond behavioral responses to light, which may affect zooplankton clearance rates in the dark versus light, differences in feeding approach and digestion likely influence whether zooplankton provide protection or enhancement. Both species use cilia to generate flow and direct particles toward them, but their feeding processes differ substantially. *T. pyriformis* use phagocytosis to capture larger particles and pinocytosis for smaller ones^56–58^. While viruses have been shown in the food vacuoles of *Tetrahymena spp*.,^5,10,18^ evidence is mixed as to whether viruses are inactivated in the food vacuole^5,10,18^. *Tetrahymena spp*. possess digestive enzymes, but the specific enzymes and their roles in viral inactivation are not known.

*B. calyciflorus* uses a ciliated corona and buccal funnel and mechanically processes food in the mastax before enzymatic digestion in the gut. For *B. calyciflorus*, viruses are likely too small to be mechanically processed, so inactivation can occur only through enzymatic digestion. *B calyciflorus* can regulate particle ingestion and reject particles^49^, but the viral particles may be too small to be selectively rejected.

Comparing the two organisms, *T. pyriformis* is likely better able to capture and process viruses, but their digestive residence times also differ. Full digestive cycles in *T. pyriformis* have been reported to last up to 2 hours^57^, with egestion of undigestible material sometimes occurring as early as 30 min ^57,59^. Another study showed that MS2 was inactivated within *T. thermophilia*, a closely related species to *T. pyriformis*, 24 hours after co-incubation^18^. The gut passage time in *B. calyciflorus* ranges from 15 to 45 minutes^60,61^. The digestive residence time could lend to physical shielding of the virus when inside the zooplankton. UV resistance of the zooplankton organisms^10^ could allow for shielding the virus from solar UV. Further studies are needed to link the digestive process and ingestion behavior to viral inactivation.

## 4. Environmental Significance

This study demonstrates that zooplankton can either act synergistically with or impede sunlight inactivation of viruses, with outcomes that depend on zooplankton and virus type, as well as on environmental context. If zooplankton provide protection, they could act as vectors that transport infectious viruses and promote viral persistence. Conversely, if zooplankton enhances decay under sunlight, they could contribute to attenuation and be considered beneficial in lowering viral risk. In complex, multi-species communities, ecological interactions such as predation and trophic transfer can shift the balance between protection and enhanced inactivation. For example, predation of *T. pyriformis* by *B. calyciflorus* can alter exposure pathways and thereby change viral fate^12,61,62^. Disinfection models should incorporate potential zooplankton contributions, but inclusion depends on a mechanistic understanding to predict whether zooplankton will protect viruses or promote their inactivation. Our results provide initial evidence of these contrasting roles, and targeted experiments are needed to identify biological processes and environmental drivers that determine zooplankton modulation of viral fate.

## Acknowledgments

We want to thank the following individuals for project support: Htet Kyi Winn for general project support; Nils Rädecker and the Meibom lab for use of equipment and lab space, Enio Zanchetta and the Ludwig lab for algae culturing support; and Pierre Rossi for training and use of equipment at CEMBL. MVO was supported by a Swiss Government Excellence Fellowship. This research was supported by the SNSF Grant IZSEZ0_214651. Images in the TOC art were obtained from Integration and Application Network (ian.umces.edu/media-library).

## Disclaimer

The authors declare no competing interests.

## Data Availability statement

All data needed to evaluate the findings and conclusions are presented in the paper, in the supporting information, and in the Zenodo repository: https://doi.org/10.5281/zenodo.18816930

## Author Contribution statement

M. I. Verbel-Olarte: Investigation, writing-original draft; T. Kohn: Formal analysis and writing-reviewing, and editing; N. Ismail: Conceptualization, supervision, investigation, formal analysis, and writing-original draft, reviewing and editing.

## TOC Art

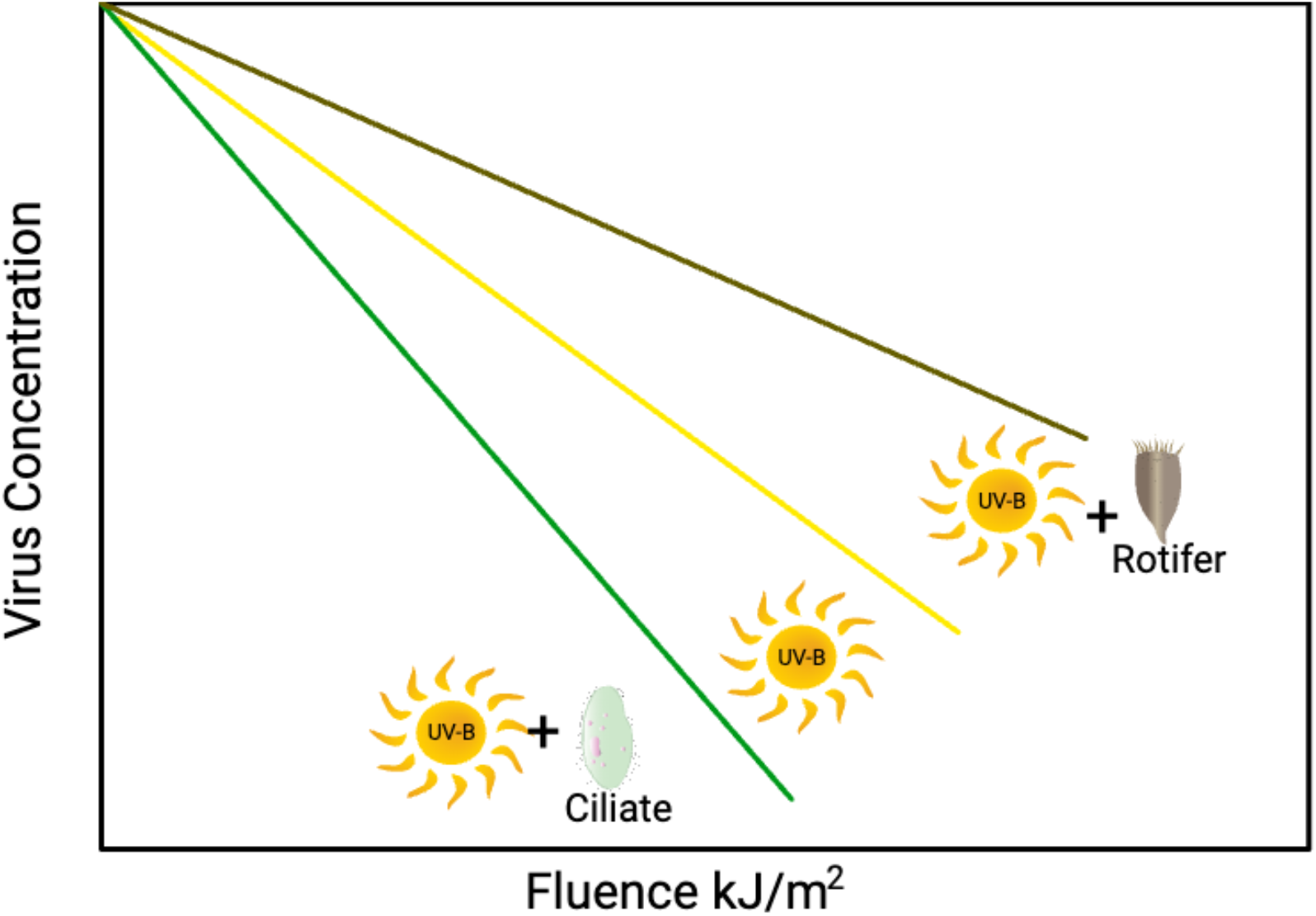

## References

(1) Declerck, S. A. J.; de Senerpont Domis, L.N. Contribution of Freshwater Metazooplankton to Aquatic Ecosystem Services: An Overview. Hydrobiologia 2023, 850 (12), 2795–2810. 10.1007/s10750-022-05001-9.

(2) Deng, L.; Krauss, S.; Feichtmayer, J.; Hofmann, R.; Arndt, H.; Griebler, C. Grazing of Heterotrophic Flagellates on Viruses Is Driven by Feeding Behaviour. Environmental Microbiology Reports 2014, 6 (4), 325–330. 10.1111/1758-2229.12119.

(3) Pinheiro, M. D. O.; Power, M. E.; Butler, B. J.; Dayeh, V. R.; Slawson, R.; Lee, L. E. J.; Lynn, D. H.; Bols, N. C. Use of Tetrahymena thermophila to Study the Role of Protozoa in Inactivation of Viruses in Water. Applied and Environmental Microbiology 2007, 73 (2), 643–649. 10.1128/AEM.02363-06.

(4) Atanasova, N. D.; Dey, R.; Scott, C.; Li, Q.; Pang, X.-L.; Ashbolt, N. J. Persistence of Infectious Enterovirus within Free-Living Amoebae–A Novel Waterborne Risk Pathway? Water Res. 2018, 144 (Journal Article), 204–214.

(5) Olive, M.; Daraspe, J.; Genoud, C.; Kohn, T. Uptake without Inactivation of Human Adenovirus Type 2 by Tetrahymena pyriformis Ciliates. Environmental Science: Processes and Impacts 2023, 25 (7), 1181–1192. 10.1039/d3em00116d.

(6) Zhang, M.; Altan-Bonnet, N.; Shen, Y.; Shuai, D. Waterborne Human Pathogenic Viruses in Complex Microbial Communities: Environmental Implication on Virus Infectivity, Persistence, and Disinfection. Environ. Sci. Technol. 2022, 56 (9), 5381–5389. 10.1021/acs.est.2c00233.

(7) Olive, M.; Moerman, F.; Fernandez-Cassi, X.; Altermatt, F.; Kohn, T. Removal of Waterborne Viruses by Tetrahymena pyriformis Is Virus-Specific and Coincides with Changes in Protist Swimming Speed. Environmental Science & Technology 2022, 56 (7), 4062–4070.

(8) Alotaibi, M. A. Internalisation of Enteric Viruses by Acanthamoeba castellanii, via Ingestion of Virus-Infected Mammalian Cells. Food and Environmental Virology 2011, 3 (3–4), 109–114.

(9) Folkins, M.; Dey, R.; Ashbolt, N. J. Interactions between Human Reovirus and Free-Living Amoebae: Implications for Enteric Virus Disinfection and Aquatic Persistence. Environ.Sci.Technol. 2020, 54 (16), 10201–10206.

(10) Akunyili, A. A.; Alfatlawi, M.; Upadhyaya, B.; Rhoads, L. S.; Eichelberger, H.; Van Bell, C. T. Ingestion without Inactivation of Bacteriophages by Tetrahymena. J.Eukaryot.Microbiol. 2008, 55 (3), 207–213.

(11) Abbas, M. D.; Nazir, J.; Stumpf, P.; Marschang, R. E. Role of Water Fleas (Daphnia magna) in the Accumulation of Avian Influenza Viruses from the Surrounding Water. Intervirology 2012, 55 (5), 365–371.

(12) Ismail, N. S.; Olive, M.; Fernandez-cassi, X.; Bachmann, V.; Kohn, T. Viral Transfer and Inactivation through Zooplankton Trophic Interactions. Environmental Science & Technology 2020, 54, 9418–9426. 10.1021/acs.est.0c02545.

(13) Benyahya, M.; Laveran, H.; Bohatier, J.; Senaud, J.; Ettayebi, M. Interactions between the Ciliated Protozoan Tetrahymena pyriformis and the Simian Rotavirus SA11. European Journal of Protistology 1997, 33 (2), 211–213. 10.1016/S0932-4739(97)80038-7.

(14) Frada, M. J.; Schatz, D.; Farstey, V.; Ossolinski, J. E.; Sabanay, H.; Ben-Dor, S.; Koren, I.; Vardi, A. Zooplankton May Serve as Transmission Vectors for Viruses Infecting Algal Blooms in the Ocean. Current Biology 2014, 24 (21), 2592–2597.

(15) Mayers, K. M. J.; Lawrence, J.; Sandnes Skaar, K.; Töpper, J. P.; Petelenz, E.; Rydningen Saltvedt, M.; Sandaa, R. A.; Larsen, A.; Bratbak, G.; Ray, J. L. Removal of Large Viruses and Their Dispersal through Fecal Pellets of the Appendicularian Oikopleura dioica during Emiliania huxleyi Bloom Conditions. Limnology and Oceanography 2021, 66 (11), 3963– 3975. 10.1002/lno.11935.

(16) Gerba, C. P. Virus Occurrence and Survival in the Environmental Waters. Perspectives in Medical Virology 2007, 17, 91–108.

(17) Olive, M.; Gan, C.; Carratalà, A.; Kohn, T. Control of Waterborne Human Viruses by Indigenous Bacteria and Protists Is Influenced by Temperature, Virus Type, and Microbial Species. Appl.Environ.Microbiol. 2020, 86 (3).

(18) Pinheiro, M. D.; Power, M. E.; Butler, B. J.; Dayeh, V. R.; Slawson, R.; Lee, L. E.; Lynn, D. H.; Bols, N. C. Inactivation of the Bacteriophage MS2 by the Ciliated Protozoan, Tetrahymena thermophila. Water Quality Research Journal 2008, 43 (1), 69–76.

(19) Verani, M.; Di Giuseppe, G.; Tammaro, C.; Carducci, A. Investigating the Role of Acanthamoeba polyphaga in Protecting Human Adenovirus from Water Disinfection Treatment. Eur.J.Protistol. 2016, 54 (Journal Article), 11–18.

(20) Gibson, K. E. Viral Pathogens in Water: Occurrence, Public Health Impact, and Available Control Strategies. Current Opinion in Virology 2014, 4, 50–57. 10.1016/j.coviro.2013.12.005.

(21) Lanrewaju, A. A.; Enitan-Folami, A. M.; Sabiu, S.; Edokpayi, J. N.; Swalaha, F. M. Global Public Health Implications of Human Exposure to Viral Contaminated Water. Front. Microbiol. 2022, 13. 10.3389/fmicb.2022.981896.

(22) DeFlorio-Barker, S.; Wing, C.; Jones, R. M.; Dorevitch, S. Estimate of Incidence and Cost of Recreational Waterborne Illness on United States Surface Waters. Environmental Health 2018, 17 (1), 3.

(23) Nelson, K. L.; Boehm, A. B.; Davies-Colley, R. J.; Dodd, M. C.; Kohn, T.; Linden, K. G.; Liu, Y.; Maraccini, P. A.; McNeill, K.; Mitch, W. A. Sunlight-Mediated Inactivation of Health-Relevant Microorganisms in Water: A Review of Mechanisms and Modeling Approaches. Environmental Science: Processes & Impacts 2018, 20 (8), 1089–1122.

(24) Gerba, C. P. Applied and Theoretical Aspects of Virus Adsorption to Surfaces. Advances in applied microbiology 1984, 30, 133–168.

(25) Bertrand, I.; Schijven, J. F.; Sánchez, G.; Wyn-Jones, P.; Ottoson, J.; Morin, T.; Muscillo, M.; Verani, M.; Nasser, A.; de Roda Husman, A.M. The Impact of Temperature on the Inactivation of Enteric Viruses in Food and Water: A Review. Journal of Applied Microbiology 2012, 112 (6), 1059–1074.

(26) Moresco, V.; Damazo, N. A.; Barardi, C. R. M. Thermal and Temporal Stability on the Enteric Viruses Infectivity in Surface Freshwater. Water Science and Technology: Water Supply 2016, 16 (3), 620–627.

(27) McMinn, B. R.; Rhodes, E. R.; Huff, E. M.; Korajkic, A. Decay of Infectious Adenovirus and Coliphages in Freshwater Habitats Is Differentially Affected by Ambient Sunlight and the Presence of Indigenous Protozoa Communities. Virology Journal 2020, 17 (1), 1–11.

(28) Li, C.; Sylvestre, É.; Fernandez-Cassi, X.; Julian, T. R.; Kohn, T. Waterborne Virus Transport and the Associated Risks in a Large Lake. Water Research 2023, 229, 119437. 10.1016/j.watres.2022.119437.

(29) Gerba, C. P.; Betancourt, W. Q. Viral Aggregation: Impact on Virus Behavior in the Environment. Environ Sci Technol 2017, 51 (13), 7318–7325. 10.1021/acs.est.6b05835.

(30) Wu, X.; Feng, Z.; Yuan, B.; Zhou, Z.; Li, F.; Sun, W. Effects of Solution Chemistry on the Sunlight Inactivation of Particles-Associated Viruses MS2. Colloids and Surfaces B: Biointerfaces 2018, 162, 179–185. 10.1016/j.colsurfb.2017.11.056.

(31) Bichai, F.; Payment, P.; Barbeau, B. Protection of Waterborne Pathogens by Higher Organisms in Drinking Water: A Review. Can.J.Microbiol. 2008, 54 (7), 509–524.

(32) Bichai, F.; Barbeau, B.; Payment, P. Protection against UV Disinfection of E. coli Bacteria and B. subtilis Spores Ingested by C. elegans Nematodes. Water Res. 2009, 43 (14), 3397– 3406.

(33) Bichai, F.; Hijnen, W.; Baars, E.; Rosielle, M.; Dullemont, Y.; Barbeau, B. Preliminary Study on the Occurrence and Risk Arising from Bacteria Internalized in Zooplankton in Drinking Water. Water science and technology 2011, 63 (1), 108–114.

(34) King, C. H.; Shotts, E. B.; Wooley, R. E.; Porter, K. G. Survival of Coliforms and Bacterial Pathogens within Protozoa during Chlorination. Appl.Environ.Microbiol. 1988, 54 (12), 3023–3033.

(35) Snelling, W. J.; McKenna, J. P.; Lecky, D. M.; Dooley, J. S. G. Survival of Campylobacter jejuni in Waterborne Protozoa. Appl Environ Microbiol 2005, 71 (9), 5560–5571. 10.1128/AEM.71.9.5560-5571.2005.

(36) Perera, I. U.; Fujiyoshi, S.; Nishiuchi, Y.; Nakai, T.; Maruyama, F. Zooplankton Act as Cruise Ships Promoting the Survival and Pathogenicity of Pathogenic Bacteria. Microbiology and Immunology 2022, 66 (12), 564–578. 10.1111/1348-0421.13029.

(37) Wang, J. A.; Aryal, O.; Brownstein, L. N.; Shwwa, H.; Rickard, A. L.; Stephens, A. E.; Lanzarini-Lopes, M.; Ismail, N. S. Zooplankton Protect Viruses from Sunlight Disinfection. Appl Environ Microbiol 2025, 91 (4). 10.1128/aem.02540-24.

(38) Li, J.; Yan, D.; Chen, L.; Zhang, Y.; Song, Y.; Zhu, S.; Ji, T.; Zhou, W.; Gan, F.; Wang, X.; Hong, M.; Guan, L.; Shi, Y.; Wu, G.; Xu, W. Multiple Genotypes of Echovirus 11 Circulated in Mainland China between 1994 and 2017. Scientific Reports 2019, 9 (1), 1–8. 10.1038/s41598-019-46870-w.

(39) Giammanco, G. M.; Filizzolo, C.; Pizzo, M.; Sanfilippo, G. L.; Cacioppo, F.; Bonura, F.; Fontana, S.; Buttinelli, G.; Stefanelli, P.; De Grazia, S. Detection of Echovirus 11 Lineage 1 in Wastewater Samples in Sicily. Science of the Total Environment 2024, 918 (February), 170519. 10.1016/j.scitotenv.2024.170519.

(40) Maurya, R.; Pandey, A. K. Importance of Protozoa Tetrahymena in Toxicological Studies: A Review. Science of The Total Environment 2020, 741, 140058.

(41) Dahms, H.-U.; Hagiwara, A.; Lee, J.-S. Ecotoxicology, Ecophysiology, and Mechanistic Studies with Rotifers. Aquatic Toxicology 2011, 101 (1), 1–12. 10.1016/j.aquatox.2010.09.006.

(42) Danes, L.; Cerva, L.; Poliovirus and Echovirus Survival in Tetrahymena pyriformis Culture In Vivo. Journal of Hygiene, Epidemiology, Microbiology, and Immunology 1984, 28, 193– 200.

(43) US Environmental Protection Agency. Methods for Measuring the Acute Toxicity of Effluents and Receiving Waters to Freshwater and Marine Organisms; US Environmental Protection Agency, Office of Water: Washington, DC, 2002.

(44) Carratalà, A.; Shim, H.; Zhong, Q.; Bachmann, V.; Jensen, J. D.; Kohn, T. Experimental Adaptation of Human Echovirus 11 to Ultraviolet Radiation Leads to Resistance to Disinfection and Ribavirin. Virus Evolution 2017, 3 (2), 1–11. 10.1093/ve/vex035.

(45) Pecson, B. M.; Martin, L. V.; Kohn, T. Quantitative PCR for Determining the Infectivity of Bacteriophage MS2 upon Inactivation by Heat, UV-B Radiation, and Singlet Oxygen: Advantages and Limitations of an Enzymatic Treatment to Reduce False-Positive Results. Applied and Environmental Microbiology 2009, 75 (17), 5544–5554. 10.1128/AEM.00425-09.

(46) Water Environment Federation and American Public Health Association. Standard Methods for the Examination of Water and Wastewater, 21st ed.; American Public Health Association (APHA): Washington, DC, USA, 2005.

(47) Silverman, A. I.; Peterson, B. M.; Boehm, A. B.; McNeill, K.; Nelson, K. L. Sunlight Inactivation of Human Viruses and Bacteriophages in Coastal Waters Containing Natural Photosensitizers. Environ.Sci.Technol. 2013, 47 (4), 1870–1878.

(48) Starkweather, P. L.; Gilbert, J. J.; Frost, T. M. Bacterial Feeding by the Rotifer Brachionus calyciflorus: Clearance and Ingestion Rates, Behavior and Population Dynamics. Oecologia 1979, 44 (1), 26–30.

(49) Gilbert, J. J.; Starkweather, P. L. Feeding in the Rotifer Brachionus calyciflorus: I. Regulatory Mechanisms. Oecologia 1977, 28 (2), 125–131. 10.1007/BF00345247.

(50) Lavin, D. P.; Hatzis, C.; Srienc, F.; Fredrickson, A. G. Size Effects on the Uptake of Particles by Populations of Tetrahymena pyriformis Cells. The Journal of Protozoology 1990, 37 (3), 157–163. 10.1111/j.1550-7408.1990.tb01120.x.1.

(51) Bischel, H. N.; Schertenleib, A.; Fumasoli, A.; Udert, K. M.; Kohn, T. Inactivation Kinetics and Mechanisms of Viral and Bacterial Pathogen Surrogates during Urine Nitrification. Environmental Science: Water Research & Technology 2015, 1 (1), 65–76.

(52) Kim, H.-J.; Lee, J.-S.; Hagiwara, A. Phototactic Behavior of Live Food Rotifer Brachionus plicatilis Species Complex and Its Significance in Larviculture: A Review. Aquaculture 2018, 497, 253–259. 10.1016/j.aquaculture.2018.07.070.

(53) Colangeli, P.; Schlägel, U. E.; Obertegger, U.; Petermann, J. S.; Tiedemann, R.; Weithoff, G. Negative Phototactic Response to UVR in Three Cosmopolitan Rotifers: A Video Analysis Approach. Hydrobiologia 2019, 844 (1), 43–54. 10.1007/s10750-018-3801-y.

(54) Kim, D. H.; Casale, D.; K\Hohidai, L.; Kim, M. J. Galvanotactic and Phototactic Control of Tetrahymena pyriformis as a Microfluidic Workhorse. Applied Physics Letters 2009, 94 (16).

(55) Rainville, J. A Search for Light-Detecting Proteins in the Free-Living Protist, Tetrahymena thermophila: Does Tetrahymena Have Opsin-like or Bacteriorhodopsin-like Proteins? Senior Honors Projects 2013.

(56) Nilsson, J. R. Fine Structure and RNA Synthesis of Tetrahymena during Cytochalasin B Inhibition of Phagocytosis. Journal of Cell Science 1977, 27 (1), 115–126.

(57) Nilsson, J. R. On Food Vacuoles in Tetrahymena pyriformis GL. The Journal of Protozoology 1977, 24 (4), 502–507. 10.1111/j.1550-7408.1977.tb01000.x.

(58) Nilsson, J. R.; Deurs, B. V. Coated Pits and Pinocytosis in Tetrahymena. Journal of Cell Science 1983, 63 (1), 209–222.

(59) Blum, J. J.; Greenside, H. Particle Ejection from the Cytoproct of Tetrahymena. The Journal of Protozoology 1976, 23 (4), 500–502. 10.1111/j.1550-7408.1976.tb03827.x.

(60) Starkweather, P. L. Aspects of the Feeding Behavior and Trophic Ecology of Suspension-Feeding Rotifers. In Rotatoria; Dumont, H. J., Green, J., Eds.; Springer Netherlands: Dordrecht, 1980; pp 63–72. 10.1007/978-94-009-9209-2_13.

(61) Joaquim-Justo, C.; Detry, C.; Caufman, F.; Thomé, J.-P. Feeding of Planktonic Rotifers on Ciliates: A Method Using Natural Ciliate Assemblages Labelled with Fluorescent Microparticles. Journal of plankton research 2004, 26 (11), 1289–1299.

(62) Mohr, S.; Adrian, R. Functional Responses of the Rotifers Brachionus calyciflorus and Brachionus rubens Feeding on Armored and Unarmored Ciliates. Limnology & Oceanography 2000, 45 (5), 1175–1179. 10.4319/lo.2000.45.5.1175.

