## Supplemental Text, Tables, and Figures for "Differential Effects of Zooplankton on Sunlight Inactivation of Viruses"

### *Chlorella vulgaris* Culture and Preparation

The freshwater microalgae *Chlorella vulgaris,* which were used as the primary food source for *B. calyciflorus*, were kindly provided by the Ludwig Group at EPFL. Individual colonies were selected from the agar plate stocks and batch-cultured in 100 mL of sterile mineral medium (Table S3) [1–3] in 250 mL Erlenmeyer flasks with cotton wool plugs. The Erlenmeyer flasks were shaken at 130 rpm on a platform shaker (C10, New Brunswick Scientific) and illuminated from below at 100 ± 5 μmol photons $m^{-2}$ $s^{-1}$on an 18:6 hour light: dark cycle.

**Table S1.** Phosphate buffer saline (PBS) using Milli-Q Water

| **Reagents** | **Chemical formula** | **Concentration [g $L^{-1}$]** | **Producer** |
|---|---|---|---|
| disodium hydrogen phosphate | $Na_2HPO_4$ | 0.71 | Acros organics |
| sodium chloride | NaCl | 0.58 | Acros organics |

Solution was autoclaved to sterilize and was adjusted to pH 7.4

**Table S2.** Moderately hard synthetic freshwater (MHSFW) using Milli-Q Water

| **Reagents** | **Chemical formula** | **Concentration [mg $L^{-1}$]** | **Producer** |
|---|---|---|---|
| sodium bicarbonate | $NaHCO_3$ | 96 | Acros organics |
| calcium sulfate dihydrate* | $CaSO_4 \cdot 2H_2O$ | 60 | AppliChem |
| magnesium sulfate heptahydrate | $MgSO_4 \cdot 7H_2O$ | 122.865 | AppliChem |
| potassium chloride | KCl | 4 | AppliChem |

*This reagent was mixed separately for 24 h on a magnetic stirrer.

After mixing all the chemicals, the MHSFW was aerated vigorously for 48 h

**Table S3:** Mineral medium preparation for microalgae growth using Milli-Q water

| Reagents | Chemical formula | Concentration [mg $L^{-1}$] | Producer |
|---|---|---|---|
| sodium nitrate | $NaNO_3$ | 1557 | Sigma-Aldrich |
| monopotassium phosphate | $KH_2PO_4$ | 118.5 | Sigma-Aldrich |
| magnesium sulfate heptahydrate | $MgSO_4 \cdot 7H_2O$ | 102 | Sigma-Aldrich |
| | EDTA-FeNa | 20 | Sigma-Aldrich |
| calcium chloride hexahydrate | $CaCl_2 \cdot 6H_2O$ | 86.8 | Sigma-Aldrich |
| boric acid | $H_3BO_3$ | 0.415 | Sigma-Aldrich |
| copper (II) sulfate pentahydrate | $CuSO_4 \cdot 5H_2O$ | 0.475 | Sigma-Aldrich |
| manganese (II) chloride tetrahydrate | $MnCl_2 \cdot 4H_2O$ | 1.65 | Sigma-Aldrich |
| cobalt(II) sulfate heptahydrate | $CoSO_4 \cdot 7H_2O$ | 0.3 | Sigma-Aldrich |
| zinc sulfate heptahydrate | $ZnSO_4 \cdot 7H_2O$ | 1.35 | Sigma-Aldrich |
| ammonium heptamolybdate tetrahydrate | $(NH_4)_6 Mo_7 O_{24} \cdot 4H_2O$ | 0.085 | Sigma-Aldrich |
| ammonium metavanadate | $NH_4VO_3$ | 0.007 | Sigma-Aldrich |

The pH was adjusted to 7.00 ± 0.05.

**Table S4.** Virus inactivation rate constant ratios based on time (k, $hr^{-1}$) and fluence ( κ, $m^2$ $kJ^{-1}$) under light and dark conditions

| **k or *κ*- value Ratio (Experimental/Control)** | ***T pyriformis*** | ***B. calyciflorus*** |
|---|---|---|
| Dark E11 ($k_{exp}/k_{control}$) | 1.8 | 3.8 |
| Light E11 ($\kappa_{exp}/\kappa_{control}$) | 3.1 | 0.57 |
| Dark MS2 ($k_{exp}/k_{control}$) | 0.66 | 2.3 |
| Light MS2 ($\kappa_{exp}/\kappa_{control}$) | 3.2 | 1.3 |

**Table S5.** Additive and observed light inactivation rate constants (k, $hr^{-1}$)

| Virus | $k_{\text{Dark, } T.\ pyriformis} + k_{\text{Light, virus-only control}}$ | $k_{\text{light, } T.\ pyriformis}$ |
|---|---|---|
| MS2 | 0.11 | 0.15 |
| E11 | 0.30 | 0.37 |

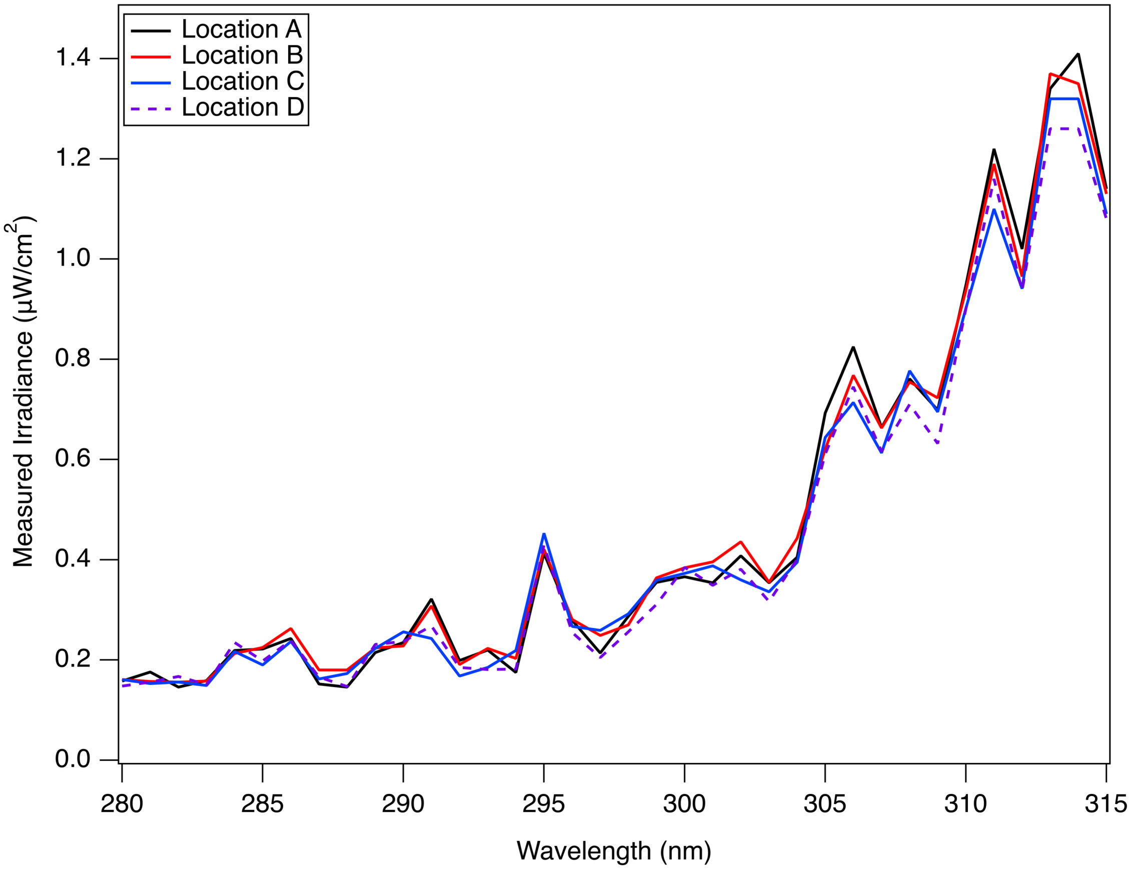


**Figure S1.** Irradiance of the 4 locations of vessels under the solar simulator for the UVB wavelength range (280-315 nm).

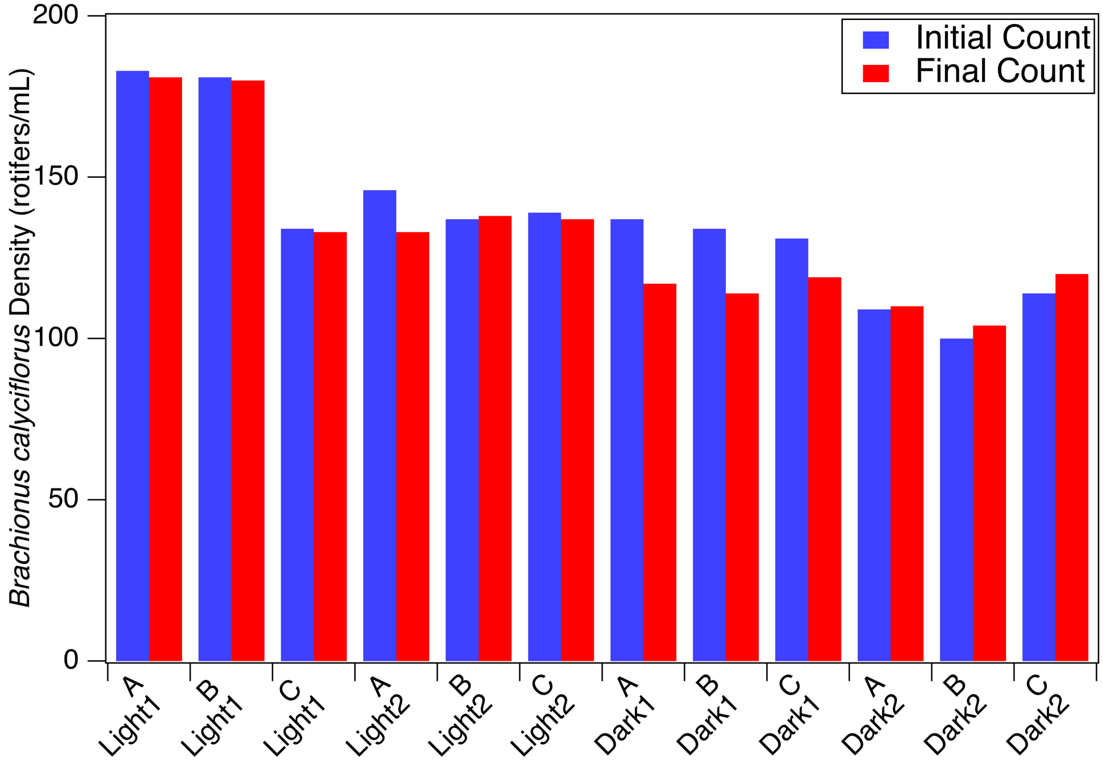


**Figure S2.** *Brachionus calyciflorus* density at the start and completion of dark and light experiments. The letters A, B, and C represent experimental replicates for each experiment type and correspond to the location beneath the solar simulator. Light represents sunlight exposure experiments, with the numbers 1 and 2 representing the first and second experiments. Dark represents the dark experiments, with numbers 1 and 2 representing the first and second experiments.

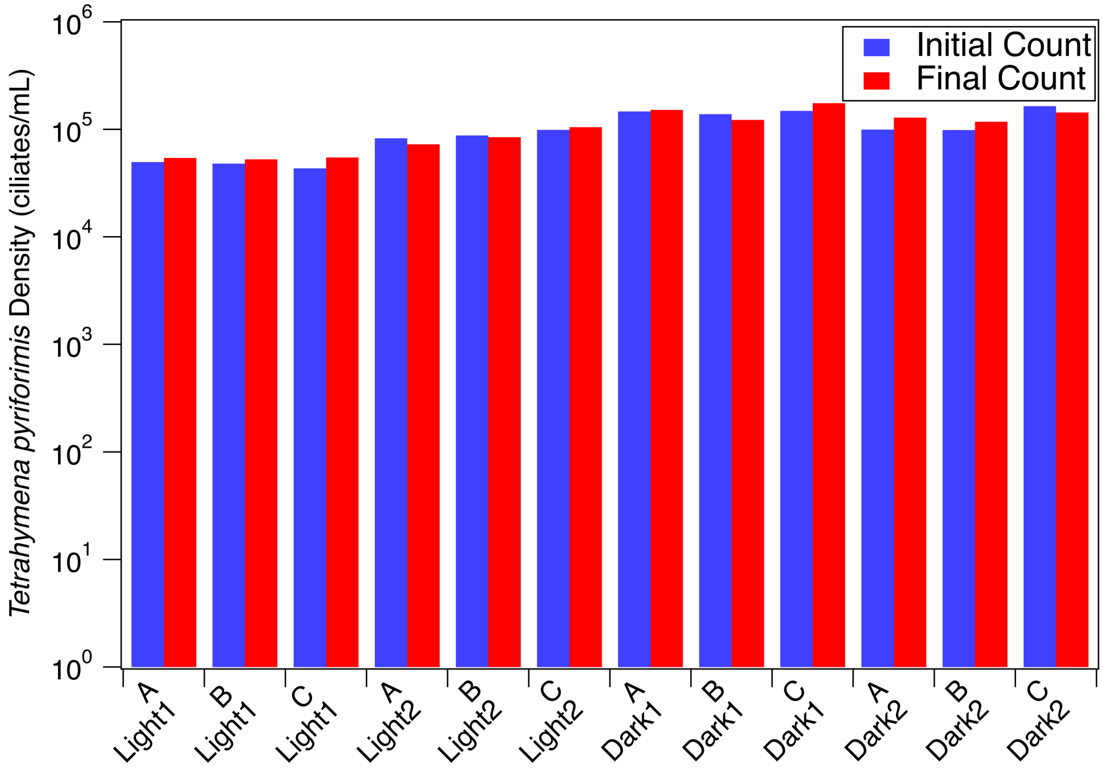


**Figure S3.** *Tetrahymena pyriformis* density at the start and completion of each experiment. The letters A, B, and C represent experimental replicates for each experiment type. Light represents sunlight exposure experiments, with the numbers 1 and 2 representing the first and second experiments. Dark represents the dark experiments, with numbers 1 and 2 representing the first and second experiments.

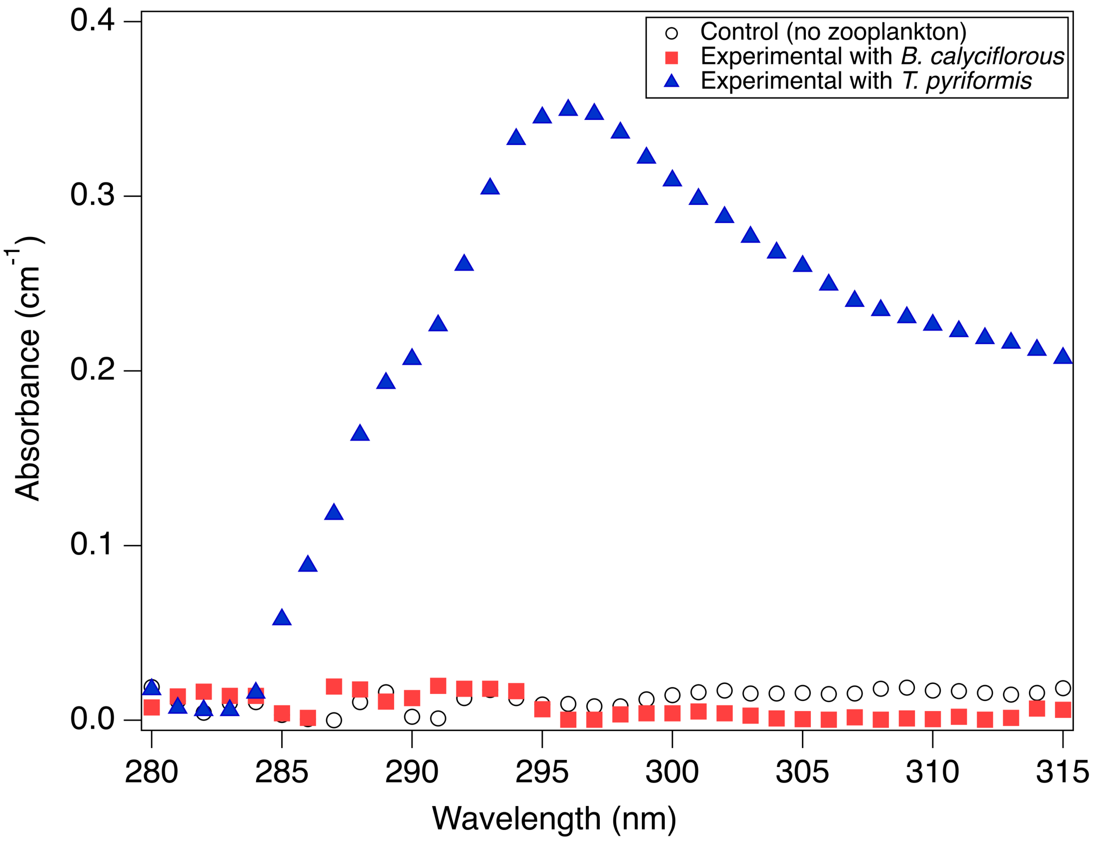


**Figure S4.** Average absorbance values for moderately hard synthetic freshwater with *T. pyriformis*, *B. calyciflorous,* and the control without zooplankton

### REFERENCES


(1) Pulgarin, A.; Giannakis, S.; Pulgarin, C.; Ludwig, C.; Refardt, D. A Novel Proposition for a Citrate-Modified Photo-Fenton Process against Bacterial Contamination of Microalgae Cultures. *Applied Catalysis B: Environmental* **2020**, *265* (October 2019), 118615. https://doi.org/10.1016/j.apcatb.2020.118615.

(2) Zanchetta, E.; Ollivier, M.; Taing, N.; Damergi, E.; Agarwal, A.; Ludwig, C.; Pick, H. Abiotic Stress Approaches for Enhancing Cellulose and Chitin Production in *Chlorella vulgaris*. *International Journal of Biological Macromolecules* **2025**, *309*, 142969.

(3) Doušková, I.; Kaštánek, F.; Maléterová, Y.; Kaštánek, P.; Doucha, J.; Zachleder, V. Utilization of Distillery Stillage for Energy Generation and Concurrent Production of Valuable Microalgal Biomass in the Sequence: Biogas-Cogeneration-Microalgae-Products. *Energy Conversion and Management* **2010**, *51* (3), 606–611. https://doi.org/10.1016/j.enconman.2009.11.008.
